# Context-dependent regulation of RpoS-dependent and RpoS-independent gene transcription by Crl during exponential growth in *Escherichia coli*

**DOI:** 10.64898/2026.10.12.758518

**Authors:** Pingzhuang Ge, Karin Schnetz, Remus T. Dame

## Abstract

RpoS, an alternative sigma factor, directly regulates approximately 10% of gene transcription in *E. coli*. The small protein Crl functions as a partner that assists RpoS in exerting its regulatory role, particularly during the stationary phase. However, the coordinating role of Crl during the exponential phase remains unclear. In addition, whether Crl can regulate gene transcription independently of RpoS is still unknown. Therefore, in this study, we used the RpoS-dependent operon *hdeAB* and the RpoS-independent operon *proVWX* as model systems to investigate the transcriptional regulatory function of Crl using RT-qPCR and 3C-qPCR. Since the expression of both Crl and RpoS is modulated by environmental signals, we further examined the transcription of these operons under different pH and osmotic conditions. The results showed that Crl can regulate transcription independently of RpoS and exhibits environment-specific bidirectional regulation. Overall, this study improves our understanding of the regulatory mechanisms of Crl and highlights the complexity of transcriptional regulation.

## Introduction

Precise transcriptional regulation underlies bacterial adaptation to fluctuating environments. In *Escherichia coli*, transcription is mediated by the RNA polymerase core enzyme composed of five subunits (α2, β, β′, ω) (1–3). Promoter-specific recognition is achieved through association of the core enzyme with sigma (σ) factors, forming the RNA polymerase holoenzyme and enabling selective initiation of gene expression (4–6). Seven sigma factors have been identified in *E. coli* (7). Typically, individual sigma factors direct transcription of distinct gene regulons (7, 8), yet some genes can be controlled by multiple sigma factors. For example, acid fitness island genes are co-regulated by the housekeeping sigma factor RpoD (σ⁷⁰) (9–12) and the general stress sigma factor RpoS (σ^38^) (9, 13–18). RpoD and RpoS are highly similar in both sequence and structure (19). Both proteins contain the conserved domains σ² and σ⁴ (19, 20). In RpoD, these domains recognize the −10 and −35 promoter elements, respectively (21–23), whereas RpoS-dependent promoters often possess weakly conserved −35 elements and rely more strongly on other promoter characteristics, such as discriminator sequences, promoter architecture, and DNA structural properties, to facilitate transcription initiation (13, 24–28). Transcription of RpoD dependent genes occurs throughout the entire bacterial life cycle (29). RpoS has a low intracellular concentration during the exponential phase. As cells enter the stationary phase or encounter other stress conditions, RpoS accumulates and reprograms transcription. RpoS regulates the expression of ∼ 10% of genes (either up or down) involved in pH response, osmotic stress, and oxidative stress resistance in *E. coli* (15, 24, 30, 31). Despite its low abundance during the exponential phase, RpoS continues to regulate the transcription of part of its target genes (7, 15, 30, 32, 33), including 8 of the acid resistance operons (*gadA*, *gadX*, *gadW*, *gadE*, *mdtE*, *hdeD*, *hdeAB* and *slp-dctR*) which are located in acid fitness islands (12 genes in total) (15).

The levels of RpoS are regulated via multiple mechanisms controlling transcription of *rpoS*, translation, and proteolytic degradation. RpoS activity can be further modulated by accessory factors that influence its association with RNA polymerase and promoter recognition. These mechanisms have been extensively reviewed in the literature (24, 34). Here, we specifically focus on the 16.5 kDa Sigma factor-binding protein, Crl. Expression of Crl protein is increased at the from exponential to stationary phase growth (35–37). Crl function depends on RpoS, structural and biochemical studies have demonstrated that Crl directly interacts with the σ^2^ domain of RpoS and the RNAP subunit β′ to stabilize the holoenzyme (20, 38). Studies by Typas et al. demonstrated that Crl acts as a fine-tuner of RpoS activity during stationary phase, facilitating the reorganization of the transcription machinery towards stress-responsive transcription. Crl not only enhances RpoS-dependent transcription but also influences sigma factor competition, thereby affecting allocation of the RNA polymerase core enzyme to RpoD- and RpoS-dependent transcriptional programs (39). Crl has also been implicated in indirect negative regulation: the RpoS-mediated silencing of *bgl* operon requires Crl (31). Previous *in vitro* studies demonstrated that Crl enhances the binding and open complex formation of RpoS– RNAP at the *hdeA* promoter (14). GFP reporter fusion assays further showed that transcription driven by the *hdeA* promoter during the stationary phase in *E. coli* is significantly reduced in the absence of either Crl or RpoS (14). Microarray analyses revealed that, when *E. coli* was grown to the exponential phase in minimal medium, the transcription level of *hdeA* was significantly reduced in the *rpoS* deletion strain compared with the wild-type strain (15, 40), indicating that Crl-dependent modulation of RpoS activity may also contribute to transcriptional regulation during exponential phase. However, the mechanism by which Crl assists RpoS in regulating gene transcription during the exponential phase remains unclear. In addition, genome-wide and promoter-reporter analyses revealed that Crl does not globally regulate the entire RpoS regulon, but instead selectively affects a subset of RpoS-dependent genes in a condition-dependent manner (14, 35). For example, the regulatory function of Crl can be influenced by the temperature at which bacteria are grown. Compared with cultures grown at 37 °C, the degree of stimulation of Crl at *glgS* and *gadB* was markedly reduced at 30 °C (14). On the other hand, the transcription of the *csgBA* operon was activated by Crl at 30 °C, but not at 37 °C (35).

In this study, we employed the RpoS-dependent operon *hdeAB* (9, 13, 40, 41) and the RpoS-independent operon *proVWX* (42, 43) as models to investigate the regulatory functions of Crl during exponential growth in *E. coli*. Specifically, we sought to determine whether Crl can influence transcription independently of RpoS, whether its regulatory effects are modulated by environmental conditions such as pH and osmolarity, and whether Crl-dependent transcriptional regulation is accompanied by changes in local chromosomal organization. To address these questions, gene expression and chromosomal interaction frequencies were analyzed using RT-qPCR and 3C-qPCR, respectively.

## Method and materials

### Strain construction

All the *E. coli* strains used in this study were derived from *Escherichia coli* MG1655. All derivative strains were constructed by using the two-step genome editing method based on the λ-red recombinase system (44) and phage P1 transduction method (45). Strains used in the current study are listed in Table 1.

**Table 1:** Strains used in this study.

| Strain ID | Description | Construction Method | Reference |
| --- | --- | --- | --- |
| MG1655 | *Escherichia coli* K-12 |  |  |
| NT331 (*crl::IS1*) | MG1655 $\Delta endA$; *crl::IS1* | λ-red recombinase system | (46) |
| XT3124 (*crl<sup>+</sup> rpoS<sup>+</sup>*) | MG1655 $\Delta endA$; *crl<sup>+</sup>* | Phage P1 transduction | This study |
| XT3132 (*crl::IS1 $\Delta rpoS$*) | MG1655 $\Delta endA$; *crl::IS1*; *rpoS::camR* | λ-red recombinase system | (40) |
| XT3155 (*crl<sup>+</sup> $\Delta rpoS$*) | MG1655 $\Delta endA$; *crl<sup>+</sup>*; *rpoS::CamR* | λ-red recombinase system | This study |

### Culture condition

All the strains were cultured overnight on LB agar plates with the appropriate antibiotic at 37 °C in an incubator (Catalog numbers: 390-0384, VWR® INCU-Line®).

For acid resistance studies, four single colonies were taken from the agar plate and these were separately grown overnight in 2 mL M9 glycerol medium (pH=6.6, 42 mM Na_2_HPO_4_, 22 mM KH_2_PO_4_, 19 mM NH_4_Cl, 2.0 mM MgSO_4_, 0.1 mM CaCl_2_, 170 mM NaCl, 1X trace elements, 1% Bacto™ casamino acids (BD), 10 μg/mL thiamine (Sigma-Aldrich), 0.4% glycerol (PanReac Applichem) with the proper antibiotic at 37 °C in a 200 rpm shaking incubator (Eppendorf Innova® 40). The overnight culture was used to inoculate M9 glycerol medium to a starting OD₆₀₀ of 0.05. The culture was grown in an incubator at 37 °C with shaking at 200 rpm until an OD₆₀₀ of 1.0 (early exponential phase). For testing at pH 6.6, 1 mL of the suspension was kept aside for RT-qPCR, 9 mL was used for 3C-qPCR. For acid shock (pH 2.0), 37% HCl was added to adjust the pH of the M9 medium to 2.0. The bacterial suspension was then incubated at 37 °C with shaking at 200 rpm for 10 minutes.

For experiments at different osmolarity, four single colonies were taken from the agar plate and the were separately grown overnight in 2 mL low salt (LS) M9 glycerol medium (42 mM Na_2_HPO_4_, 22 mM KH_2_PO_4_, 19 mM NH_4_Cl, 2.0 mM MgSO_4_, 0.1 mM CaCl_2_, 80 mM NaCl, 1X trace elements, 1% Bacto™ casamino acids (BD), 10 μg/mL thiamine (Sigma-Aldrich), 0.4% glycerol (PanReac Applichem) with proper antibiotic at 37 °C with shaking at 200 rpm (Eppendorf Innova® 40). The overnight cultures were used to inoculate M9 glycerol medium to a starting OD₆₀₀ of 0.05. The culture was grown in an incubator at 37 °C with shaking at 200 rpm until an OD₆₀₀ of 1.0. For the low salt group, milliQ water was added to the suspension with a ratio of 46 μL per mL of culture, 1 mL of the suspension was kept aside for RT-qPCR, 9 mL was used for 3C-based studies, 10 mL used for ChIP-based study. For the salt shock group, 5M NaCl was added to the suspension with a ratio of 46 μL per mL of culture resulting in a final concentration of 300 mM NaCl, 1 mL of the suspension was kept aside for RNA isolation, 9 mL was used for 3C-based study.

### RNA isolation

All of the RNA samples were isolated by using PureLink® RNA Mini Kit (Catalog numbers: 12183020, Thermo Fisher scientific) according to the manufacturer’s protocol of purification from bacterial cells. The RNA concentration was measured by using NanoDrop™ 2000 spectrophotometer (Catalogue number: ND-2000, Thermo Scientific™) and NanoDrop 2000/2000c software version 1.6. The quality of the RNA sample was analyzed by running 1% gel-red pre-stained agarose gel.

### RT-qPCR

#### Primer design

All primers (Supplementary table1) used for RT-qPCR were designed based on the *Escherichia coli* K-12 MG1655 reference genome (Accession number: NC_000913) and were synthesized by Sigma-Aldrich (In solution, 100 μM). Primer specificity was confirmed by melting curve analysis.

#### Reaction set-up

RT-qPCR experiments were performed as described previously (46).

#### Data analysis

Data analysis was performed as described previously (46). To ensure comparability of gene expression levels, raw data are typically normalized using reference genes with relatively stable expression. According to RT-qPCR guidelines (47), at least three reference genes should be used to ensure data reliability. In this study, *dnaA*, *rpoD* and *rrsA* were used as reference genes. *dnaA* encodes the chromosomal replication initiator protein DnaA. *RpoD* encodes the RNA polymerase σ^70^ factor. *DnaA* and *rpoD* both have been used as reference control for RT-qPCR experiments in *E. coli* in different pH (48, 49) and osmolarity (46). *RrsA* codes for 16S rRNA. It has been used to normalize relative transcript levels in RT-qPCR studies of the *proVWX* operon (50). In the Results and Discussion sections of this study, all RT-qPCR data were normalized using *dnaA* as the reference control.

### 3C-qPCR

#### 3C library preparation

*E. coli* cells were harvested by centrifugation after cultivation under experimental conditions. Preparation of 3C libraries was performed as previously described (40). The restriction enzyme MluCI was used to digest the chromosome in all 3C studies of the *hdeAB* operon and its flanking regions. Library quality was assessed by electrophoresis on a 1% gel-red pre-stained agarose gel.

#### Control library preparation

A synthetic control library was prepared for 3C-PCR (Supplementary table 2). The synthetic control template was generated by individually amplifying the chimeric fragments of interest by PCR and pooling them at equimolar ratios. This control library was then serially diluted to produce the standard samples for 3C-qPCR.

#### Primer and TaqMan probe design

All primers (Supplementary table 3) used for 3C-qPCR were designed on the *Escherichia coli* K-12 MG1655 reference genome (Accession number: NC_000913.3 – https://www.ncbi.nlm.nih.gov/nuccore/556503834) and ordered from Sigma-Aldrich (In solution, 100 μM). The HPLC-purified, double-quenched TaqMan probes (Supplementary table 4) were designed as described previously (51), which is with a 5’ 6-FAM fluorophore, 3’ Iowa Black™ Fluorescence Quencher and an internal ZEN quencher were ordered from Integrated DNA Technologies. All the primers and probes were dissolved in 1X TE to a final concentration of 100 μM and stored at -20°C.

#### Reaction set-up and data analysis

The PrimeTime™ Gene Expression Master Mix Kit (Catalogue number: 10557710, Integrated DNA Technologies, Inc.) was used for 3C-qPCR. The experiment’s reaction setup and data analysis were performed as described previously (46, 52).

### Data visualization and statistical analyses

Data visualization for RT-qPCR and 3C-qPCR studies and the associated statistical analyses were performed by using GraphPad Prism (version 10.1.0). Statistical significance was determined by unpaired, two-tailed Student’s t-test. P-values indicate the level of significance: *p* < 0.05 (*), *p* < 0.01 (**), *p* < 0.001 (***) and ns indicates no significant difference.

### Genomic DNA isolation

A single *E. coli* (NT331) colony was cultured in 2 mL LB medium up to stationary phase at 37 °C with shaking at 200 rpm. Genomic DNA was isolated as previously described (46). Genomic DNA was dissolved in nuclease-free water. The DNA concentration was measured using the Qubit® dsDNA HS Assay Kit (Catalogue number: Q32854, ThermoFisher Scientific).

## Results

### Crl regulates *hdeAB* transcription in a growth condition–dependent manner during exponential growth

The *E. coli* K-12 strain NT331, routinely used as reference strain in our laboratory, carries an insertion sequence (IS1) in the *crl* gene, resulting in inactivation of Crl (53, 54). Therefore, we constructed a *crl*⁺ derivative of this strain by replacing the *crl::IS1* allele with *crl^+^*. In addition, to investigate the RpoS-independent function of Crl, we generated *crl*⁺ Δ*rpoS* strains (**Table 1**). As we observed in the previous and current study, the transcription of *hdeAB* operon and its flanking genes are induced by acid shock (40) **(Figure 1a-c)**. We further compared the transcription response of *hdeAB* and its flanking genes to acid shock in the *crl*⁺ *rpoS*⁺ strain. Although *hdeA* and *hdeD* remained inducible by acid shock, the overall transcriptional response of the *hdeAB* operon and its flanking genes were attenuated compared with the Crl-deficient strain **(Figure 1c)**, as *hdeB* no longer exhibited significant acid-induced activation **(Figure 1a)**. In our previous study, acid shock also stimulated transcription of both the *hdeAB* operon and the upstream *hdeD* gene in the *crl::IS1* Δ*rpoS* strain (40). However, in the *crl*⁺ Δ*rpoS* strain, only *hdeA* and *hdeD* remained acid inducible, while the transcription level of *hdeB* was unaffected by acid shock **(Figure 1b)**. These results suggest that Crl represses the acid-responsive induction of the *hdeAB* operon during exponential growth. When the *crl::IS1 rpoS*⁺ strain was compared with the *crl*⁺ *rpoS*⁺ strain, restoration of *crl* was found to decrease transcription of the *hdeAB* operon and the upstream *hdeD* gene during exponential growth (**Figure 1d**), indicating a repressive effect of Crl on these genes during the exponential phase. This observation contrasts with previous studies showing that Crl promotes *hdeA* transcription during stationary phase, when cellular RpoS levels are substantially elevated (14, 55). These findings suggest that the regulatory effect of Crl on *hdeA* is growth phase dependent and may reflect differences in both RpoS abundance and Crl activity across growth phases. To further investigate the regulatory roles of Crl and RpoS, we examined the transcriptional profile of *csgB*, a gene known to be directly regulated by both RpoS and Crl (35, 56). The results showed that *csgB* transcription was markedly reduced in the *crl::IS1* Δ*rpoS* strain compared with *crl::IS1 rpoS*^+^ strain (**Figure 1e**), which is consistent with previous studies. Interestingly, in the Crl-restored strain, deletion of *rpoS* did not alter the transcription level of *csgB* (**Figure 1e**), consistent with the transcriptional pattern observed for *hdeD* (**Figure 1f**). In the *crl*⁺ strains, mutation of *rpoS* decreased the transcription of the *hdeAB* operon (**Figure 1f**), which is also in agreement with previous reports (15, 40). Compared with the *crl::IS1 rpoS*^+^, in the *crl*^+^ *rpoS*^+^ strain, restoring *crl* did not affect the transcription level of *csgB* in exponential phase (**Figure 1e**), but in *rpoS* deletion strains, the transcription of *csgB* was activated when *crl* was restored (**Figure 1e**).

**Figure 1:**
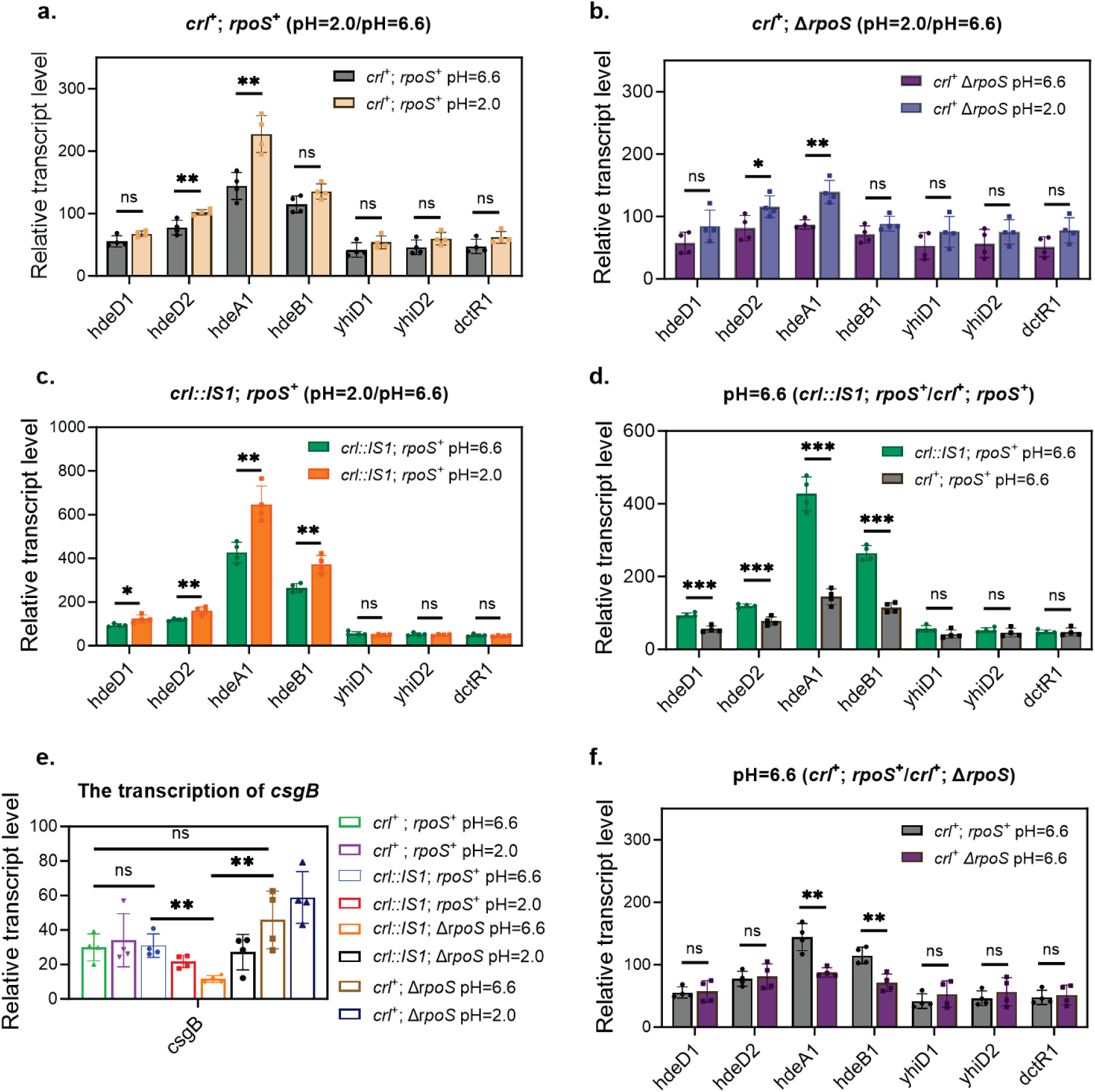
Transcription profile of the *hdeAB* operon and its flanking genes at pH 6.6 and pH 2.0 conditions in the *crl*^+^ *rpoS*^+^. (**a**), *crl*^+^ Δ*rpoS* (**b**), *crl::IS1 rpoS*^+^ (**c**). Transcription profile of *hdeAB* operon and its flanking genes at pH 6.6 in the *crl*^+^ *rpoS*^+^ strain compared with *crl::IS1 rpoS*^+^ strain (**d**). Transcription profile of *csgB* at pH 6.6 and pH 2.0 in different strains (**e**). Transcription profile of *hdeAB* operon and its flanking genes at pH 6.6 in the *crl*^+^ *rpoS*^+^ strains compared with *crl*^+^ Δ*rpoS* (**e**). Bar graphs (follow left y-axis) represent relative transcript levels. Each group includes four biological replicates and three technical replicates. *dnaA*, *rpoD* and *rrsA* were tested as internal reference genes for normalization, yielding similar results; *dnaA* is used for data presentation. Statistical significance was determined by unpaired, two-tailed Student’s t-test. *p*-values indicate the level of significance: *p* < 0.05 (*), *p* < 0.01 (**), *p* < 0.001 (***) and *ns* indicates no significant difference.

Previous studies have revealed that the effect of Crl on RpoS-dependent transcription is temperature-dependent, with Crl generally exerting a stronger positive effect on expression of a subset of RpoS-regulated genes at low temperatures (14). To investigate the influence of growth conditions on the regulatory function of Crl, we also examined the transcription of *hdeAB* under low salt (0.08M NaCl) and salt shock (0.08M to 0.3M NaCl) conditions. First, the transcription of *hdeAB* was not affected by salt shock **(Figure 2a-c)**. Although a significant decrease in *hdeA* transcription was observed in the *crl::IS1 rpoS*^+^ strain following salt shock, this result may have been influenced by a single outlier data point (**Figure 2c**). In addition, the previously observed repression of *hdeAB* transcription by Crl at 0.17 M NaCl condition (pH 6.6 group) **(Figure 1d)** disappeared under low salt conditions (0.08M NaCl) **(Figure 2d)**. These results indicate that the transcriptional regulatory activity of Crl is salt concentration– dependent. In strains with active Crl, deletion of RpoS did not result in any detectable change in *hdeAB* transcription **(Figure 2e)**. Transcription in these strains is driven by RpoD. Furthermore, in the RpoS-deficient background, complementation of Crl also did not alter the transcription level of *hdeAB* **(Figure 2f)**. This indicates that RpoD driven transcription of *hdeAB* is not affected by Crl, as expected.

**Figure 2:**
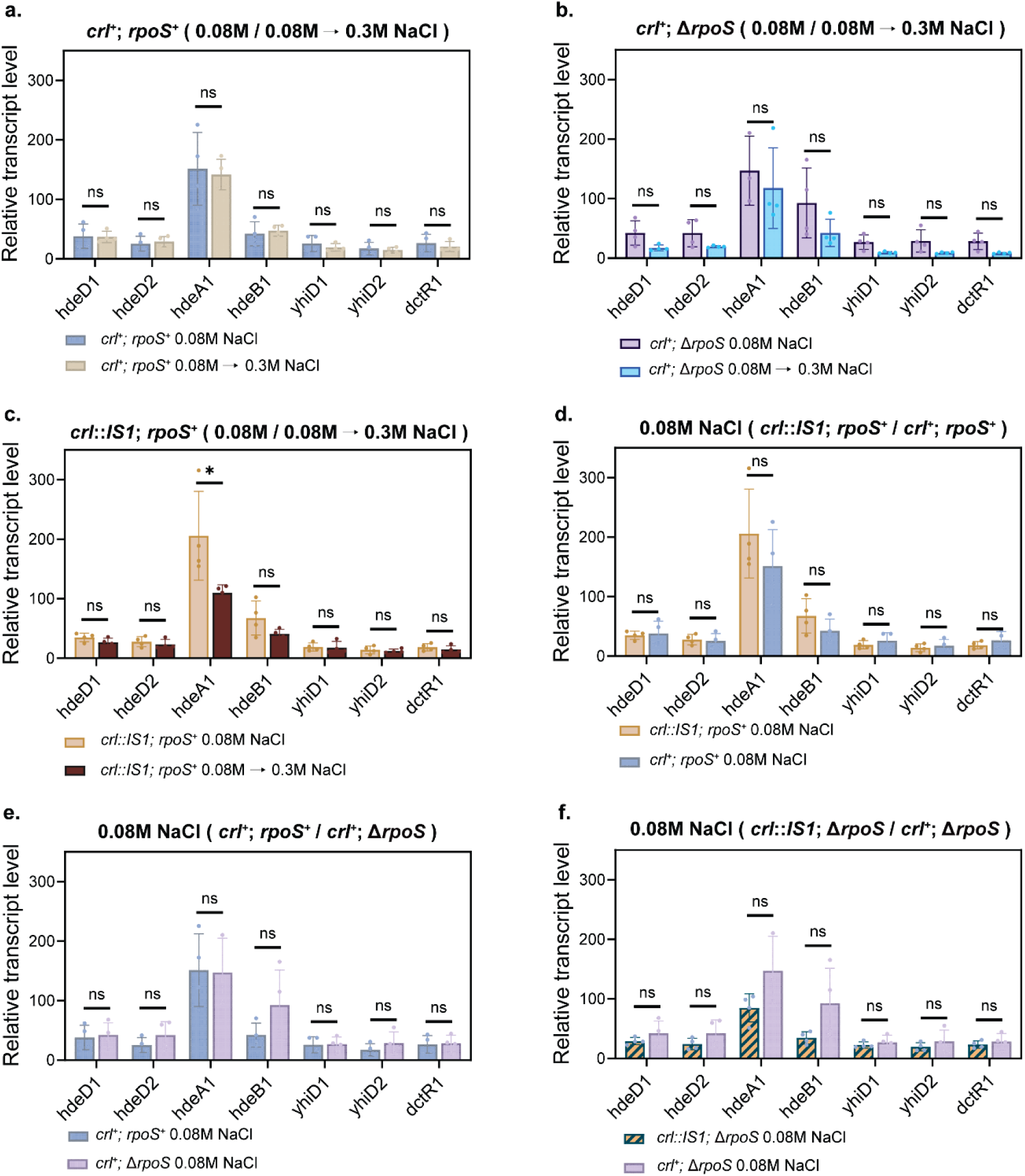
Transcription profile of the *hdeAB* operon and its flanking genes at low salt (0.08M NaCl) and salt shock (0.08M→0.3M NaCl) conditions in the *crl*^+^ *rpoS*^+^. (**a**), *crl*^+^ Δ*rpoS* (**b**), *crl::IS1 rpoS*^+^ (**c**). Transcription profile of *hdeAB* operon and its flanking genes at low salt (0.08M NaCl) in the *crl*^+^ *rpoS*^+^ strain compared with *crl::IS1 rpoS*^+^ strain (**d**). Transcription profile of *hdeAB* operon and its flanking genes at low salt (0.08M NaCl) in the *crl*^+^ *rpoS*^+^ (**e**) *crl::IS1* Δ*rpoS* (**f**) strains compared with *crl*^+^ Δ*rpoS*. Bar graphs (follow left y-axis) represent relative transcript levels. Each group includes four biological replicates and three technical replicates. *dnaA*, *rpoD* and *rrsA* were tested as internal reference genes for normalization, yielding similar results; *dnaA* is used for data presentation. Statistical significance was determined by unpaired, two-tailed Student’s t-test. *p*-values indicate the level of significance: *p* < 0.05 (*), *p* < 0.01 (**), *p* < 0.001 (***) and *ns* indicates no significant difference.

### Crl does not affect the local chromatin structure of the *hdeAB* operon

The histone-like nucleoid structuring protein (H-NS) functions as a transcriptional repressor of the *hdeAB* operon (40, 57, 58). The transcription activation of *hdeAB* at acidic conditions is attributed to decreased H-NS occupancy at the *hdeAB* promoter, leading to loss of the H-NS-mediated hairpin-like chromatin structure (40). To investigate whether Crl affects the local chromosomal architecture of the *hdeAB* locus, we performed 3C-qPCR analysis in the *crl::IS1 rpoS*^+^, *crl*^+^ *rpoS*^+^ and *crl*^+^ Δ*rpoS* strains. Briefly, an anchor fragment encompassing the *hdeAB* promoter region (Fragment 10, color in red) was used to quantify its interaction frequency with a series of neighboring test fragments spanning the locus (**Figure 3**). Compared with *crl::IS1 rpoS*^+^ strain, restoration of *crl* did not affect local chromatin structure **(Figure 3a)**, indicating that the Crl-mediated transcription repression of *hdeAB* was not through altered local chromatin structure. The relative interaction frequency between fragment 8/9/12 and anchor fragment (fragment 10) was decreased after acid shock in both *crl*^+^ *rpoS*^+^ and *crl*^+^ Δ*rpoS* strains **(Figure 3b, c)** which is consistent with our previous results (40). Interestingly, compared with the *crl*^+^ *rpoS*^+^ strain, the relative interaction between fragment 9 and the anchor fragment was decreased in the *crl*^+^ Δ*rpoS* strain **(Figure 3d)**, but the biological meaning (and cause) remains unclear.

**Figure 3.**
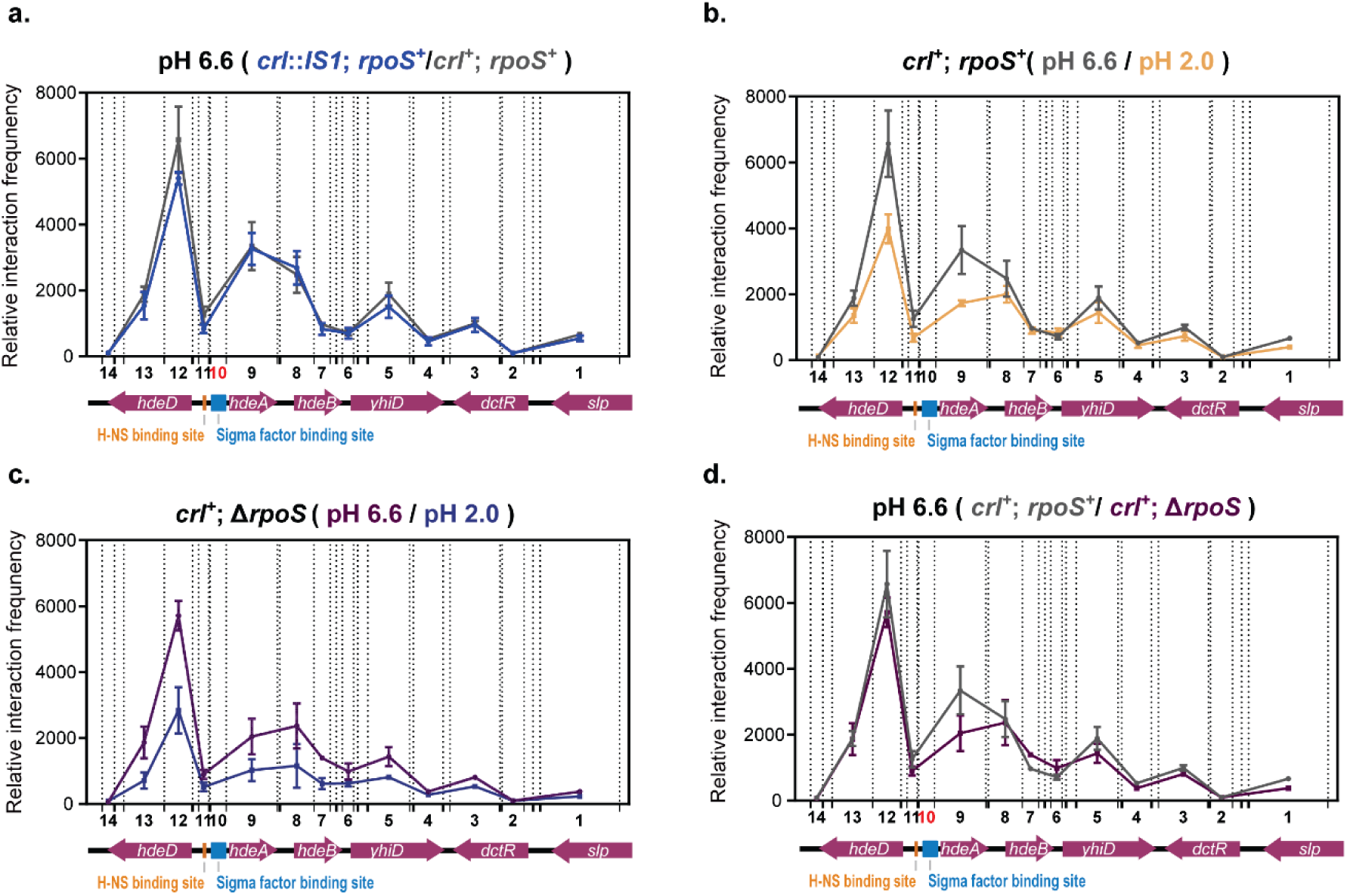
Local chromatin architecture of the *hdeAB* operon and its flanking genes. Relative interaction frequency of the *hdeAB* operon and its flanking genes at pH 6.6 condition in *crl*^+^ *rpoS*^+^ strains compared with *crl::IS1 rpoS*^+^ (**a**). Relative interaction frequency at the *hdeAB* operon and its flanking genes in *crl*^+^ *rpoS*^+^ (**b**), *crl*^+^ Δ*rpoS* (**c**) strains at pH 6.6 and pH 2.0. Relative interaction frequency of the *hdeAB* operon and its flanking genes at pH 6.6 condition in *crl*^+^ Δ*rpoS* strains compared with *crl*^+^ *rpoS*^+^ (**d**). Each group includes four biological replicates and three technical replicates. Dashed lines indicate the MluCI restriction enzyme cleavage sites within the *hdeAB* operon. The x-axis shows the relative positions and lengths of target fragments and the anchor fragment (fragment 10, highlighted in red) used in this study. The y-axis represents the relative interaction frequency between the anchor and target fragments.

### Crl functions in the regulation of RpoS-independent gene transcription

As an RpoS-independent operon (42, 43), we also tested the gene transcription profile of *proVWX* in *crl*⁺ *rpoS*^+^, *crl*⁺ Δ*rpoS* and *crl::IS1 rpoS*^+^ strains in different pH conditions. Although several *proVWX* transcripts displayed statistically significant differences following acid shock, the observed changes were relatively modest and did not reveal a consistent acid-induced transcriptional response across the operon (**Figure 4a-c**). Restoring *crl*, the transcription of *proVWX* increased by around 10% at pH 6.6 **(Figure 4d)**, suggesting that Crl influences expression of this RpoS-independent operon. Whether this effect is mediated directly by Crl or indirectly through secondary regulatory pathways remains unclear. We also examined *proVWX* transcription under different osmotic conditions. Since *proVWX* is an osmotically responsive operon, as expected, its transcription levels increased markedly following salt shock (**Figure 5a-c**). However, the observed positive regulatory effect of Crl on *proVWX* transcription **(Figure 4d**) was not detected under low salt conditions **(Figure 5d**). More interestingly, under salt shock conditions, Crl negatively impacted *proVWX* transcription **(Figure 5e**). These findings indicate that the effect of *Crl* somehow differs depending on salt concentration. When comparing the *crl::IS1* Δ*rpoS* strain with the *crl*⁺ Δ*rpoS* strain, we found that under low-salt conditions, Crl again upregulated the transcription of *proV* and *proW* **(Figure 5f**), whereas no effect of Crl was evident under conditions of salt shock **(Figure 5g**).

**Figure 4:**
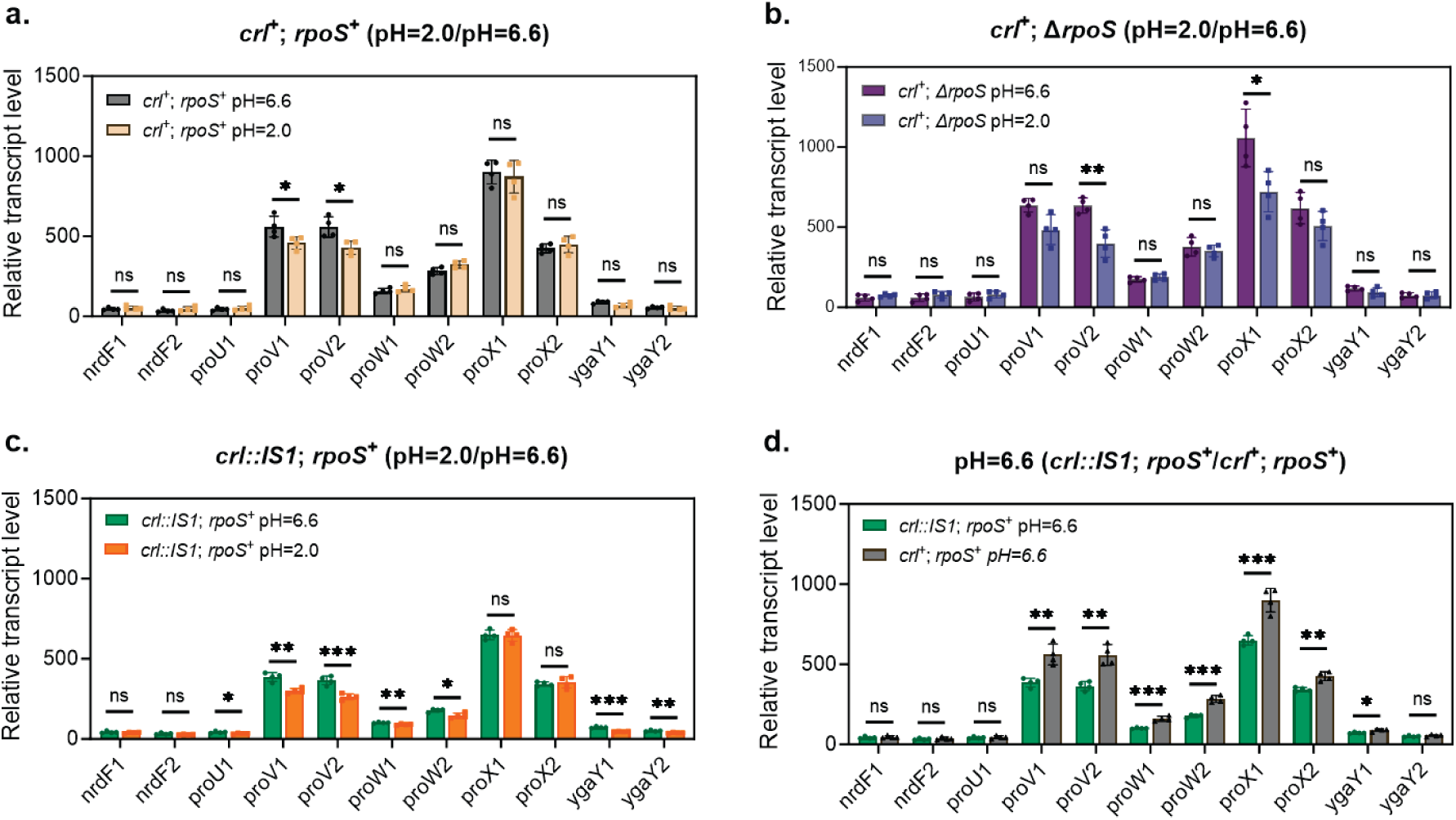
Transcription profile of the *proVWX* operon and its flanking genes at pH 6.6 and pH 2.0 conditions in the *crl*^+^ *rpoS*^+^. (**a**), *crl*^+^ Δ*rpoS* (**b**), *crl::IS1 rpoS*^+^ (**c**). Transcription profile of *proVWX* operon and its flanking genes at pH 6.6 in the *crl*^+^ *rpoS*^+^ strain compared with *crl::IS1 rpoS*^+^ strain (**d**). Bar graphs (follow left y-axis) represent relative transcript levels. Each group includes four biological replicates and three technical replicates. *dnaA*, *rpoD,* and *rrsA* were tested as internal reference genes for normalization, yielding similar results; *dnaA* is used for data presentation. Statistical significance was determined by unpaired, two-tailed Student’s t-test. *p*-values indicate the level of significance: *p* < 0.05 (*), *p* < 0.01 (**), *p* < 0.001 (***) and *ns* indicates no significant difference.

**Figure 5:**
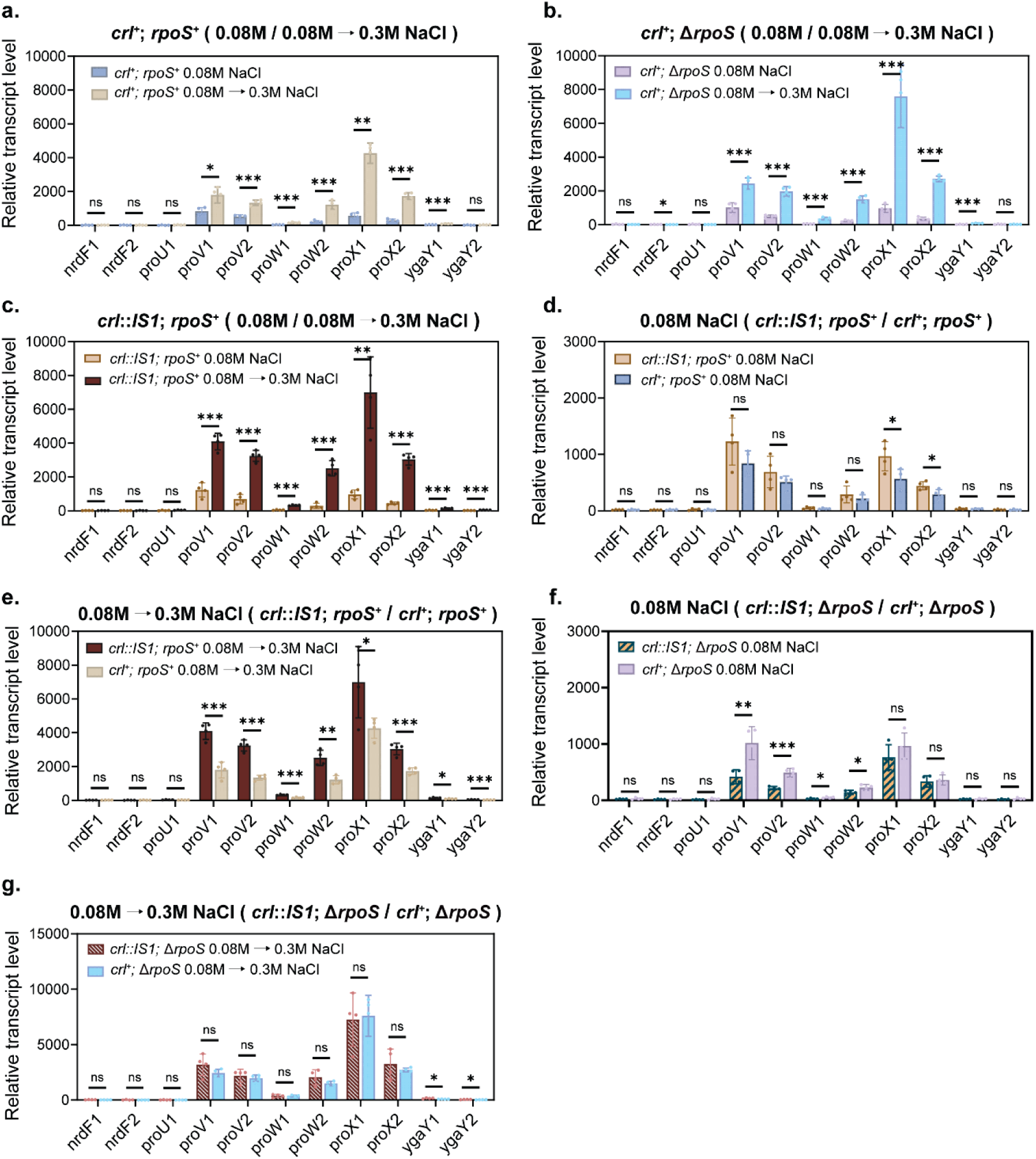
Transcription profile of the *proVWX* operon and its flanking genes at low salt (0.08M NaCl) and salt shock (0.08M→0.3M NaCl) conditions in the *crl*^+^ *rpoS*^+^. (**a**), *crl*^+^ Δ*rpoS* (**b**), *crl::IS1 rpoS*^+^ (**c**). Transcription profile of *proVWX* operon and its flanking genes at low salt (0.08M NaCl) (**d**) or salt shock condition (0.08M→0.3M NaCl) (**e**) in the *crl*^+^ *rpoS*^+^ strain compared with *crl::IS1 rpoS*^+^ strain. Transcription profile of *proVWX* operon and its flanking genes at low salt (0.08M NaCl) (**f**) and salt shock (0.08M→0.3M NaCl) (**g**) in the *crl::IS1* Δ*rpoS* strain compared with *crl*^+^ Δ*rpoS*. Bar graphs (follow left y-axis) represent relative transcript levels. Each group includes four biological replicates and three technical replicates. *dnaA*, *rpoD* and *rrsA* were tested as internal reference genes for normalization, yielding similar results; *dnaA* is used for data presentation. Statistical significance was determined by unpaired, two-tailed Student’s t-test. *p*-values indicate the level of significance: *p* < 0.05 (*), *p* < 0.01 (**), *p* < 0.001 (***) and *ns* indicates no significant difference.

## Discussion

A critical step in bacterial transcription initiation is the association of a sigma factor with RNA polymerase (RNAP) to form the holoenzyme (4, 59). In *E. coli*, seven sigma factors are present, which must compete for the limited pool of RNAP core enzyme (60). Among them, RpoD serves as the primary housekeeping sigma factor and controls the transcription of most genes (4, 59). In contrast, RpoS is the key sigma factor governing gene expression during the stationary phase and in response to various stress conditions (24). To ensure proper adaptation of the transcriptional program to changing environments, RpoS expression is tightly controlled at the transcriptional, translational, and post-translational levels (24, 34, 61). Previous studies have shown that, compared with other sigma factors, RpoS exhibits the lowest binding affinity for RNAP (60). Crl has been reported to stimulate transcription by promoting the assembly of the RpoS–RNAP complex and facilitating the formation of the RNA polymerase–promoter open complex (RPo) (20, 38, 62).

However, the functional role of Crl remains incompletely understood. For example, it is still unclear whether Crl can influence gene transcription in the absence of RpoS and whether its regulatory activity depends on environmental conditions or growth phase. Previous studies have shown that Crl promotes RpoS-dependent *hdeA* expression during stationary phase using a promoter-reporter system (14), whereas our RT-qPCR analysis of the native chromosomal locus revealed a repressive effect of Crl on *hdeA* transcription during exponential growth (**Figure 1d**). The opposite effects of Crl on *hdeA* transcription during exponential and stationary phase may reflect differences in the regulatory environment of the promoter. In particular, cellular ppGpp levels increase substantially upon entry into stationary phase and are known to influence transcription in a promoter-specific manner (63). Methodological differences between promoter-reporter assays and measurements of endogenous transcript levels should also be considered. Reporter constructs may not fully recapitulate the chromosomal context, local DNA topology, or higher-order regulatory mechanisms acting at the native *hdeAB* locus. Therefore, future studies employing RNA-seq, RT-qPCR, or other transcriptome-based approaches to directly quantify gene expression from native chromosomal loci will help contribute to a more comprehensive understanding of Crl-mediated gene regulation. Under both low-salt and salt shock conditions, Crl did not exhibit detectable regulatory effects on *hdeA* transcription (**Figure 2d**), indicating that Crl function is also environment-dependent. A similar phenomenon was observed in the transcriptional regulation of the RpoS-independent *proVWX* operon. Specifically, at 0.17 M NaCl, pH 6.6, Crl exhibited a positive regulatory effect on *proVWX* transcription (**Figure 4d**). In contrast, under low-salt conditions (0.08 M NaCl), Crl showed no detectable regulatory effect on the operon (**Figure 5d**), whereas under salt shock conditions, Crl clearly repressed *proVWX* transcription (**Figure 5e**). Both Crl and RpoS accumulate progressively during bacterial growth and reach peak levels in the stationary phase (15, 35). In addition, RpoS expression is induced by environmental stimuli such as acid stress and increased osmolarity (24, 34, 40), while Crl expression can also be activated at low temperature (35). Therefore, changes in the intracellular levels of Crl and/or RpoS represent one possible explanation for the differential regulation of *hdeA* transcription observed under different growth phases and environmental conditions. Quantifying protein levels under different environmental conditions is essential for understanding the dynamic regulatory mechanisms of Crl and RpoS. Global proteomic studies in *E. coli* have shown that several post-translational modifications, including lysine acetylation (64–67), phosphorylation (68–70) and succinylation (71), vary substantially with growth phase, nutrient availability and metabolic state. Although post-translational regulation of Crl has not been reported, such mechanisms could potentially contribute to the condition-dependent effects of Crl observed in this study.

In addition to Crl and RpoS, H-NS can regulate the transcription of *hdeAB* operon by remodeling the hairpin-like structure within the promoter region (40, 57). Our experimental results demonstrate that the presence of Crl does not affect the H-NS-mediated chromatin architecture, suggesting that different regulatory pathways controlling *hdeAB* transcription may function independently to some extent. However, this conclusion is limited to the exponential phase. During the stationary phase, when RpoS levels are high, insertion of active Crl may lead to detectable changes in chromatin structure. This hypothesis is supported by previous atomic force microscopy (AFM) studies showing that H-NS cannot repress transcription mediated by the RpoS–RNAP holoenzyme (9).

The diverse effects of Crl observed for both *hdeA* and *proVWX* suggest that Crl-dependent regulation is highly context-dependent and may not solely reflect direct activation of target promoters. By promoting formation of the RpoS–RNAP holoenzyme, Crl could alter sigma factor competition and redistribute RNA polymerase between RpoS - and RpoD-dependent transcriptional programs, thereby generating both positive and negative transcriptional effects. In addition, previous studies have shown that different promoters exhibit distinct degrees of Crl dependence (14, 39), indicating that promoter architecture and transcriptional kinetics may strongly influence the outcome of Crl regulation. Although Crl has been shown to interact directly with both RpoS and the β′ subunit of RNAP (20, 38), current evidence suggests that productive activation requires RpoS, consistent with structural studies demonstrating that the major Crl-interaction surface present in RpoS is not conserved in RpoD (20). Together, these observations support a model in which the transcriptional effects of Crl arise from a combination of direct RpoS-dependent activation and indirect changes in transcriptional network organization that varies across promoters and environmental conditions.

## Data availability

All data generated in this study have been deposited in the 4TU Repository (https://doi.org/10.4121/78165b92-0a9d-427e-a004-8bcf736aed1c)

## Acknowledgements

This work was supported by the Dutch Research Council [OCENW.GROOT.2019.012 to RTD], China Scholarship Council [grant no. 202103250002 to PG].

