## Supplementary tables for "Context-dependent regulation of RpoS-dependent and RpoS-independent gene transcription by Crl during exponential growth in *Escherichia coli*"

**Supplementary files**

Supplementary table1: Primers used for RT-qPCR.

| **Primers** | **Sequences (5'-3')** |
|---|---|
| hdeD_RT-qPCR_1.fwd | CCGATGGTGTCTGTAACGC |
| hdeD_RT-qPCR_1.rev | GCTTAACGAACAAACTGGCG |
| hdeD_RT-qPCR_2.fwd | GCAATCCAGTTTATTGCCGTGC |
| hdeD_RT-qPCR_2.rev | AATGCACCCACTACTGTGC |
| hdeA_RT-qPCR_1.fwd | TCGTCCACAGCCAGGAAATC |
| hdeA_RT-qPCR_1.rev | CTGCCAGTTGTGAGCAATGC |
| hdeB_RT-qPCR_1.fwd | CATGCAACCGGGGTCATTG |
| hdeB_RT-qPCR_1.rev | GCTGTAGCGGCTTTGTCACT |
| yhiD_RT-qPCR_1.fwd | CACTGAACCATAAATACCCAGTTCG |
| yhiD_RT-qPCR_1.rev | TGTAGGTCTGACGACGGCAG |
| yhiD_RT-qPCR_2.fwd | GAAGCTGGATATCAATGGCGAC |
| yhiD_RT-qPCR_2.rev | CGGTATCAATGCTCGACTGG |
| dctR_RT-qPCR_1.fwd | TAAACGCTGCCTGGGGAAAG |
| dctR_RT-qPCR_1.rev | GTAACTGCCATTGATGAGTGCGAC |
| gade_RT-qPCR_1.fwd | ACGCTCAATATTTCGCAACAAAC |
| gade_RT-qPCR_1.rev | GTGATACCCAGGGTGACGATG |
| nrdF_RT-qPCR_1.fwd | ATCGCCTGACCAGCAATTTC |
| nrdF_RT-qPCR_1.rev | CGAGCAGCGTCAGGCCAGTA |
| nrdF_RT-qPCR_2.fwd | CGCAGAAATGGCGGAAGTGAATC |
| nrdF_RT-qPCR_2.rev | TCACATAAGAGGAGCCTGAACCG |
| proU_RT-qPCR_1.fwd | CGCTATCTTTGACAAAAAATATCAACTTTCTCG |
| proU_RT-qPCR_1.rev | GAATCTGAGGCAACCCCTGATGG |
| proV_RT-PCR_1.fwd | TCCGACGCCGAACTCCG |
| proV_RT-PCR_1.rev | ACGCAGTATTGTCCAGCACG |
| proV_RT-PCR_2.fwd | GGCGAATGATTATGTCCGTACCTTCTTC |
| proV_RT-PCR_2.rev | CGAAGCCAGGGGTTTTACGAAT |
| proW_RT-PCR_1.fwd | CCATTTCCGTCCCGTCTTCCAG |
| proW_RT-PCR_1.rev | GGCGATGAGAGCGAAAACGATAATC |
| proW_RT-PCR_2.fwd | GCCAGATGCTGTTCAAAGTTCAGTTAC |
| proW_RT-PCR_2.rev | GCGATGACCACCATAGAAAGGGC |
| proX_RT-PCR_1.fwd | GAAAGATCCGAAGATCGCCAAACTGTTC |
| proX_RT-PCR_1.rev | GTTGATCGCACCTTCGCAGC |
| proX_RT-PCR_2.fwd | TATCGTTGCCAACAAAGCCTGGG |
| proX_RT-PCR_2.rev | TAATGGCGTTCTGGGCGTTAATATCTG |
| ygaY_RT-qPCR_1.fwd | CTTTTCCCTTTCCGCCAGTTCG |
| ygaY_RT-qPCR_1.rev | CAGTAAGGTCATCGAGACAATCAGG |
| ygaY_RT-qPCR_2.fwd | GCCGCTCCACCTTTTAACTACAG |
| ygaY_RT-qPCR_2.rev | GGTGTGGTGCGATTTGCCCTTAT |
| Fw_hns1_RT_qPCR | GAACAACATCCGTACTCTTCGTGC |
| Re_hns1_RT_qPCR | CTTCTAATTTTTCCAGCATTTCTTCCAGC |
| Fw_stpA1_RT-qPCR | CGAATTCTCCATTGACGTTCTTGAAG |
| Re_stpA1_RT-qPCR | CTCTGCCAGTTCACGCTGC |
| rpoD_RT-qPCR.fwd | GACGAAGAAGATGGCGATGACGAC |
| rpoD_RT-qPCR.rev | TTCCTGAGCGGTAGCGTGACTG |

| | |
|---|---|
| rrsA_qRT-PCR_fwd | CTCTTGCCATCGGATGTGCCCA |
| rrsA_qRT-PCR_rev | CCAGTGTGGCTGGTCATCCTCTCA |
| dnaA_RT-qPCR_control_fwd | CGATCTAACGTACGTGAGCTGG |
| dnaA_RTqPCR_control_rev | GCACGAAGTCGATGGTGATCG |

Supplementary table 2: Primers for standard 3C library preparation of the *hdeAB* operon

| No. | Internal name | Sequences (5'-3') |
|---|---|---|
| **1** | Fw_IC1_Probe2_1 | GTATCCGTTTTGCCATTGATAACGTTGATAACCTTCCCACCAAAGCGCGCTTGTTGACCAACATATAACCCC |
| **1** | Re_IC1_Probe2_1 | CAAAGGCAATAACCAACCTGATATTCAAAAAAGTTTTGTTGCTGTTCATAACCAGCCGGGGTTATATGTTGGTC |
| **1** | Fw_IC1_Probe2_2 | GGTTATTGCCTTTGATATTTTGCGGATTTCGACACTGAGGTTATAACCTGGTTTTCTGTATATGTCATG |
| **2** | Fw_IC2_Probe2_1 | CGTCGATCTCTTCTTCGTTCTGTATATGAACGACATTACCTTTACTCAGAATG |
| **2** | Re_IC2_Probe2_1 | GAAATACCAGGGATACGATGTTCTTCACCGCGATGAAAAACATTCTGAGTAAAGGTAATG |
| **2** | Fw_IC2_Probe2_2 | CATCGTATCCCTGGTATTTCGACACTGAGGTTATAACCTGGTTTTCTGTATATGTCATG |
| **3** | Fw_IC3_Probe2_1 | GTAACTGCCATTGATGAGTGCGACAAAATCAATGATAGTGATAGTCCGCGGAACAAAGGTCACCGGCACTTTC |
| **3** | Re_IC3_Probe2_1 | TAAAACCTGTCCATGTCATTATTTCTCCCCCTTTAATATTAAACGCTGCCTGGGGAAAGTGCCGGTGAC |
| **3** | Fw_IC3_Probe2_2 | GACATGGACAGGTTTTAATCGTTCAATTTCGACACTGAGGTTATAACCTGGTTTTCTGTATATGTCATG |
| **4** | Fw_IC4_Probe2_1 | CTGATTTAGTTTCGTTGCAGGAAAATGAAGATCATGAAGTGGTCGCCATTGATATCCAGCTTCATGCCAC |
| **4** | Re_IC4_Probe2_1 | CACCCCTGCCATACCTTTTAATAATCGCAACAGGTCTTCTATCGACGTTGTGGCATGAAGCTGGATATC |
| **4** | Fw_IC4_Probe2_2 | GGTATGGCAGGGGTGAAAGGCGTTTCAATTTCGACACTGAGGTTATAACCTGGTTTTCTGTATATGTCATG |
| **5** | Fw_IC5_Probe2_1 | GTAGGTCTGACGACGGCAGCGGATATCTGGGTGACCGCCGCCATAGGTATGGTTATTGGCAGCGGTATGTAC |
| **5** | Re_IC5_Probe2_1 | CTTCCAGCACCAACAAGGTCATCACTGAACCATAAATACCCAGTTCGTACATACCGCTGCCAATAAC |
| **5** | Fw_IC5_Probe2_2 | CTTGTTGGTGCTGGAAGTCTTCCATCAATTTCGACACTGAGGTTATAACCTGGTTTTCTGTATATGTCATG |
| **6** | Fw_IC6_Probe2_1 | GAGTAGCAAGTTGAGCCATCTTGCTGCTCCTTTTTGCATTTTTATATGACAGC |
| **6** | Re_IC6_Probe2_1 | CCAGGTTATAACCTCAGTGTCGAAATTCTGCTGTCATATAAAAATGC |
| **6** | Fw_IC6_Probe2_2 | GAATTTCGACACTGAGGTTATAACCTGGTTTTCTGTATATGTCATG |

| | | |
|---|---|---|
| **7** | Fw_IC7_Probe2_1 | GGTGGATGCTGCATGAAGAAACAGTATATAAAGGTGGCGATACCGTTACTTTAAATG |
| **7** | Re_IC7_Probe2_1 | GTTATAACCTCAGTGTCGAAATTTGAGTGAGATCGGTTTCATTTAAAGTAACGGTATCGCC |
| **7** | Fw_IC7_Probe2_2 | AATTTCGACACTGAGGTTATAACCTGGTTTTCTGTATATGTCATGTTG |
| **8** | Fw_IC8_Probe2_1 | CATCTCTCCGTAAAGCGTTTATTTTTATGGGCGCTGTAGCGGCTTTGTCACTGGTGAACG |
| **8** | Re_IC8_Probe2_1 | GTCATATCTTTAGCGGATTCATTGGCTGCCAACGCAGATTGTGCGTTCACCAGTGACAAAGCCG |
| **8** | Fw_IC8_Probe2_2 | CCGCTAAAGATATGACCTGCCAGGAATTTCGACACTGAGGTTATAACCTGGTTTTCTGTATATGTCATG |
| **9** | Fw_IC9_Probe2_1 | GTAACCCCAGCTATCGTTCAGGCTTGTACTCAGGATAAACAAGCCAACTTTAAAG |
| **9** | Re_IC9_Probe2_1 | GAAATTTTGTCCCATTCGCCTTTAACTTTATCTTTAAAGTTGGCTTGTTTATC |
| **9** | Fw_IC9_Probe2_2 | GGCGAATGGGACAAAATTTCGACACTGAGGTTATAACCTGGTTTTCTGTATATGTCATG |
| **11** | Fw_IC11_Probe2_1 | GAAGAAAATCCCCTGCTATCAATCTATGCCAAAAACGCGTCTAAG |
| **11** | Re_IC11_Probe2_1 | CCTCAGTGTCGAAATTTTTATTAAATCGACTGCATTCTTAGACGCGTTTTTGG |
| **11** | Fw_IC11_Probe2_2 | AAAATTTCGACACTGAGGTTATAACCTGGTTTTCTGTATATGTCATG |
| **12** | Fw_IC12_Probe2_1 | GATACACAGCAACCCGACGATAAACAGCAGCACGGCAATAAACTGGATTGCTCTGC |
| **12** | Re_IC12_Probe2_1 | GAAGTTTGATCTGGAGATGCTTAAAAAACATCGCAGAGCAATCCAGTTTATTG |
| **12** | Fw_IC12_Probe2_2 | CATCTCCAGATCAAACTTCAAAATTTCGACACTGAGGTTATAACCTGGTTTTCTGTATATGTCATG |
| **13** | Fw_IC13_Probe2_1 | CAAAAATGCCCAGCTCCGGTGCGCGGATGAAGAAATAGCCGATCAATAAATAG |
| **13** | Re_IC13_Probe2_1 | CTGGCCGGTATTATCCGGTTTCCTCGTCGCAGTCGCCTATTTATTGATCGGC |
| **13** | Fw_IC13_Probe2_2 | GAAACCGGATAATACCGGCCAGAAATTTCGACACTGAGGTTATAACCTGGTTTTCTGTATATGTCATG |
| **14** | Fw_IC14_Probe2_1 | GTAGCGGAGAGAGTATTAATCGGATCATAGTCACATCAAGTGAC |
| **14** | Re_IC14_Probe2_1 | GAAATTACCCCGGTTGTCACCCGGATCATAGTCACTTGATGTGACTATG |
| **14** | Fw_IC14_Probe2_2 | GACAACCGGGGTAATTTCGACACTGAGGTTATAACCTGGTTTTCTGTATATGTCATG |

| Fragment No. | Internal name | Sequences (5'-3') |
|---|---|---|
| **1** | hdeAB-yhiD1_MluCI | GTATCCGTTTTGCCATTGATAAC |
| **2** | hdeAB-yhiD2_MluCI | CGTCGATCTCTTCTTCGTTC |
| **3** | hdeAB-yhiD3_MluCI | GTAACTGCCATTGATGAGTGC |
| **4** | hdeAB-yhiD4_MluCI | CTGATTTAGTTTCGTTGCAGG |
| **5** | hdeAB-yhiD5_MluCI | GTAGGTCTGACGACGG |
| **6** | hdeAB-yhiD6_MluCI | GAGTAGCAAGTTGAGCCATC |
| **7** | hdeAB-yhiD7_MluCI | GGTGGATGCTGCATGAAG |
| **8** | hdeAB-yhiD8_MluCI | CATCTCTCCGTAAAGCG |
| **9** | hdeAB-yhiD9_MluCI | GTAACCCCAGCTATCGTTCAG |
| **10** | hdeAB-yhiD10_MluCI(Anchor primer) | GCATCTGTAACTCATTGTATTGAAA |
| **11** | hdeAB-yhiD11_MluCI | GAAGAAAATCCCCTGCTATCAATC |
| **12** | hdeAB-yhiD12_MluCI | GATACACAGCAACCCGAC |
| **13** | hdeAB-yhiD13_MluCI | CAAAAATGCCCAGCTCCG |
| **14** | hdeAB-yhiD14_MluCI | GTAGCGGAGAGAGTATTAATCG |

Supplementary table 3: Primers used for 3C-qPCR.

Supplementary table 4: The sequence of TaqMan probe for the *hdeAB* operon

| Name | Sequences (5'-3') | Purification |
|---|---|---|
| **TaqMan probe_*hdeAB*** | FAM/TGAGGTTATAACCTGGTTTTCT/ZEN/GTATATGTCATGTTGATGG/IBFQ | RP-HPLC |
